# Isolation and molecular characterization of an enteric isolate of the Genotype-I Bovine coronavirus with notable mutations in the receptor binding domain of the spike glycoprotein and deletion downstream the RNA binding domain of the nucleocapsid protein

**DOI:** 10.1101/2024.08.03.606496

**Authors:** Abid Shah, Phillip Gauger, Maged Gomaa Hemida

## Abstract

Bovine coronavirus (BCoV) continues to be a significant threat to cattle populations despite the implementation of vaccination programs. The continuous circulation of BCoV highlights the necessity for ongoing genomic surveillance to understand better the virus’s evolution and its impact on cattle health. The main goal of this study was to do isolation and perform a comprehensive molecular characterization of a new enteric field isolate of the BCoV. To identify any genetic elements in the sequences of this BCoV isolate that could act as genetic markers for BCoV infection in cattle. To achieve these objectives, the newly identified BCoV isolate was propagated on the MDBK cell line for several subsequent blind passages. The immunofluorescence assay verified confirmation of the virus propagation. We plaque purified this isolate and titrated it by plaque assay using the HRT-18 cell line. We examined the viral protein expression using the SDS-PAGE followed by the Western blot using the BCoV/S and BCoV/N and BCoV/S antibodies. Our results show a substantial increase in the viral genome copy number, protein expression, and virus infectivity of this BCoV isolate with the increase in cell culture passages. The full-length genome sequence of this isolate using the NGS was drafted. The vial genome is 31 Kb in length. The viral genome has the typical BCoV organization (5’-UTR-Gene- 1- HE-S-M-E-N-UTR-3’). Our phylogenetic analysis based on the nucleotide sequences of the (full-length genome, S, HE, and N) showed that the BCoV-13 clustered with other members of the BCoV (genotype I-i). The sequence analysis shows several synonymous mutations among various domains of the S glycoprotein, especially the receptor binding domain. We found nine notable nucleotide deletions immediately downstream of the RNA binding domain of the nucleocapsid gene. Further gene function studies are encouraged to study the function of these mutations on the BCoV molecular pathogenesis and immune regulation/evasion. This research enhances our understanding of BCoV genomics and contributes to improved diagnostic and control measures for BCoV infections in cattle populations.

## 1 Introduction

Bovine coronavirus (BCoV) is still circulating in many cattle populations throughout the world, including the USA, despite the vaccination of some of these cattle herds. BCoV was initially reported as an enteric pathogen associated with calf diarrhea by Mebus and colleagues in 1973 in a cattle herd in Missouri [1]. McNulty and colleagues isolated a respiratory form of BCoV from a cattle herd with bronchopneumonia [2]. Since then, there have been continuous reports of BCoV in cattle herds across the US [3]. Some studies showed the involvement of BCoV in the lower respiratory tract of cattle with BRDC in Iowa [4]. A recent study showed the emergence of new BCoV variants showing mutations in the R3 loop of the HE gene in samples collected from cattle herds in 12 Midwest states in the US [5]. Thus, there is a high demand for the continuous monitoring of various BCoV isolates on the genomic levels [6]. BCoV belongs to the order *Nidovirales,* Family*, Coronaviridae,* Subfamily*, Orthocoronavirinae,* Genus*: Betacoronavirus,* Subgenus*, Embecovirus* [7]. BCoV genome is a positive sense RNA about 31 Kbp in length. The BCoV genome has a typical coronavirus genome organization. The BCoV genomes are flanked by two untranslated regions at their ends. The 5’ end of the genome is occupied by a large gene called Gene-1, which is composed of two overlapping open reading frames (ORF1a and ORF1b) with a ribosomal frameshift in between. Gene-1 is cleaved by some proteases into 16 non-structural proteins (NSP-1-NSP16). At the same time, the 3’ one-third of the genome is occupied mostly by the viral structural proteins (Spike (S), Matrix (M), smell envelop (E ), and the nucleocapsid protein (N)) interspersed by some non-structural proteins. This is in addition to an additional important structural protein called Hemagglutinin (HE). The HE is present in both the BCoV and the Human coronavirus-OC43. There is also a small protein called I overlapped with the N protein. The genome organization is as follows (5’-UTR-ORF1a, ORF1b, HE, S, M, E, N, I-3-UTR). BCoV is a pneumotropic virus affecting cattle of various ages, causing several clinical syndromes (calf diarrhea (CD), winter dysentery (WD), and participated in the development of the bovine respiratory disease complex (BRDC) [6]. BCoV is endemic in most cattle populations across the globe, including in the USA [5, 8]. Despite the presence of some minor differences between the enteric and respiratory isolates of the BCoV at both the nucleotide (nt) and the amino acid (aa) levels, the viral genomes of both isolates are similar [9]. A previous study suggested that a minor alteration in one (aa) could contribute to the shift in the phenotypes of the BCoV strains [5]. Another recent study showed the correlation between some bovine genetic loci and the susceptibility of certain breeds of cattle to BCoV infection in general and the development of respiratory diseases in dairy and feedlot cattle [10]. The severity of the BCoV infection in cattle ranged from self-limiting, mild, moderate, to severe and fatal outcomes [11–13]. BCoV infection causes high economic losses to the cattle industry in the US, Japan, Canada, Europe, and South America, including high morbidity and mortality rates, especially in severe cases in young calves, decreased milk production, poor feed conversion rate, and the cost of veterinary care and biosecurity measures [14–16]. The average drop in the milk yield ranged from (0-70%) in some affected cattle herds in these areas [17–20]. Cohort studies on cattle herds in Sweden reported an average of 51-liter loss of milk per infected cow from one week before the infection until 19 days after the infection [20]. Actively shedding BCoV-infected beef cattle showed a marked reduction in body weight gain by an average of 8.17 kg per animal [21]. In Canada, a high seroprevalence of BCoV was reported among feedlot cattle in Alberta and Ontario (100% and 81%), respectively [15]. Another study showed high detection rates of BCoV-RNA in some healthy and diarrheic calves in Ontario using RT-PCR assays [22]. A recent study showed a strong association between the BCoV and the BRDC in western Canadian feedlot cattle based on metagenomic sequencing [23]. In Europe, BCoV was detected in the respiratory tract of at least 73% of cattle in 93 dairy farms [24]. Several genotypes of BCoV have been reported based on the method used for their classification and phylogenetic analysis [25]. The main goal of this study was to isolate and comprehensively characterize some newly circulating BCoV strains in cattle populations in the US by analyzing the full-length genome sequences. We intend to elucidate the phylogenetic relationships between this new isolate and existing BCoV strains in the GenBank database, and to identify potential genetic markers that could indicate BCoV infection in cattle.

## 2. Materials and Methods

### 2.1. BCoV isolates, processing of samples, and testing for bovine respiratory panel of viruses

A total of 19 samples were received from the Veterinary Diagnostic Laboratory, Iowa State University ( ISU VDL). There were 11 lung tissue suspensions and eight fecal suspensions. All samples were tested for the bovine respiratory viral panels, including (BVDV1, 2, BRSV, IBRV, bovine rotavirus, and BCoV) using the qRT-PCR assay. We selected three samples that only showed positive results for the BCoV as a sole viral agent and have a low Ct value. Initially, we tried propagating those three samples in cell culture, as shown below, but we finally focused on the BCoV-13 isolate. This isolate was collected from calves born in March 2024 from a herd of 80 crossbred beef cows in Iowa. At 3-4 days of age (this calf was three days old), the calves become lethargic and develop diarrhea. The calf died, and postmortem, the calf had pasty feces in the colon. This sample was tested negative for the other viral agents by real-time PCR as described below, mainly ( Bovine Respiratory Syncytial Virus (BRSV), Bovine Viral Diarrhea Virus (BVDV), and bovine rotavirus).

### 2.2. Isolation and propagation of the BCoV field isolate

The BCoV was isolated from a field sample as described previously [26, 27]. Briefly, the suspension from the obtained sample was centrifuged, and the supernatant was filtered through (MF-Millipore™ Membrane Filter, 0.45 µm, HAWP04700 Millipore, Sigma). The filtered supernatant was treated with 10 ug/mL of TPCK trypsin (Thermo Fisher Scientific; REF: 20233) and incubated at 37°C for 30 minutes with rocking after every 10 minutes. Subsequently, the Madine Darby Bovine Kidney (MDBK) and the Human Rectal Tumor-18 (HRT-18) cells were infected independently with the TPCK trypsin-treated filtered supernatant. The infected cells were incubated for 60 minutes at (37°C and 5% CO_2_). The inoculum was removed, and cells were washed three times with sterile PBS. The MDBK-BCoV infected cells were cultured on the Minimum Essential Medium Eagle media (Sigma-Aldrich, Cat. No. M0200-500ML). The HRT-18 cells infected with BCoV isolate were cultured using the RPMI-1640 (ATCC, 30-2001), supplemented with 10% horse serum and 1% streptomycin and penicillin antibiotics at (37°C and 5% CO_2_ ) for 3 to 5 dpi. The cells were observed daily under the inverted microscope for the development of any cytopathic effect (CPE) for up to 5 dpi. At the end of the incubation time, the cell culture supernatants and the cells were collected for further experiments.

### 2.3. Viral Plaque Assay and Plaque Purification

Cell culture supernatant from each BCoV-13 passage group was collected, frozen, and thawed three times. The supernatant was treated with 10 ug/mL of TPCK trypsin (as described earlier). The HRT-18 cells cultured in 6-well plates (about 90% confluent) were infected with 10-fold serial dilution of BCoV-13 supernatant at (37°C and 5% CO_2_) for 1 hour. Subsequently, the inoculum was removed, cell monolayer was washed with plain medium, and incubated with 3ml of media per well of 1.5% Sekam ME Agarose (Lonza; Cat. No. 50011), 2X EMEM (quality Biological; Cat. No. 115-073-101) supplemented with 1% Penicillin Streptomycin (Gibco, Ref # 15140-122) and 1 ug/ml TPCK Trypsin. The infected plates were incubated for 2 to 3 days at (37°C and 5% CO_2_). The infected cells were observed daily under the inverted microscope. We add 2-ml per well of the secondary overlay containing 1.5% Sekam ME Agarose (Lonza, Cat. # 50011), 2X EMEM (Quality Biological, Cat. # 115-073-101) supplemented with 1% Penicillin Streptomycin (Gibco, Ref. # 15140-122) and 0.33% neutral red (Thermo Scientific, Cat. # J62643.18). The plates were then incubated for 2 to 6 hours until the plaques developed. The plaques were counted and recorded per dilution. The viral titer was calculated using the Reed and Muench method as described earlier [28]. For plaque purification, we picked up several single plaques with a 10ul pipette tip and then incubated them individually in a 12-well plate containing 90% confluent HRT-18 or MDBK cells for 1 hour at (37°C and 5% CO_2_.) The inoculum was removed, and the cell monolayers were washed with plain medium three times, then cultured for 3 to 5 days with a complete medium at 37°C and 5% CO_2_ until we observed the CPE as described earlier. The plaque size was measured in mm for 10 plaques in each group by using ImageJ software [29].

### 2.4. Extraction of the viral RNAs, oligonucleotides, cDNA synthesis, and the quantitative Real Time-Polymerase Chain Reaction (qRT-PCR) analysis

The total RNAs from the original sample, the cell culture supernatants of the (HRT-18 and MDBK) cells infected with BCoV, and the sham infected (PBS) control cells were isolated using TRIzol LS Reagent (Invitrogen; Ref # 10296010) as per the manufacturer’s instructions. The concentration and quality of the extracted RNAs were measured using the NanoDrop OneC (Thermo Scientific). The primers targeting the BCoV-M gene were used to check the tested samples for the presence of the BCoV. The sequences of the BCoV-Forward primer 5’-CTGGAAGTTGGTGGAGTT-3’, while the BCoV-Reverse primer sequences 5’- ATTATCGGCCTAACATACATC-3’ [30]. The Bovine b-actin Forward primer 5’- CAAGTACCCCATTGAGCACG-3’, Bovine b-actin Reverse primer 5’- GTCATCTTCTCACGGTTGGC- 3’, were used to normalize the BCoV expression in the tested samples. The cDNAs were prepared using the high-capacity reverse transcription kit (Applied Biosystems; Lot. # 2902953) following the manufacturer’s instructions. The cDNA samples were then used to perform the qRT-PCR assay for the tested samples. The qRT-PCR was performed using the PowerUp SYBR Green Master Mix (Applied Biosystems; Lot. #2843446) following the manufacturer’s instructions. The qRT-PCR was performed using QuantStudio3 (Applied Biosystems). The BCoV viral genome expression was normalized to the bovine β-actin using the 2^−ΔΔCt method [31].

### 2.5. The immunofluorescence assay (IFA)

The MDBK cells were grown in 6-well plates and infected with BCoV-13 at the Multiplicity Of Infection (MOI) 1 for 1 hour at 37°C and 5% CO_2_. Subsequently, the cell culture supernatants were removed, and the cell monolayer was washed with the PBS three times. These BCoV-infected cells were cultured in a medium containing Minimum Essential Medium Eagle media (MEM), 10% horse serum, 1% streptomycin/penicillin solution, and culture at 37°C and 5% CO_2_ for three days. Next, the cells were fixed with 80% acetone solution (Sigma; Cat. # 179124) for 10 minutes at 37°C. These cells were washed three times with the PBS and incubated with ice-cold methanol for 10 minutes at (-20 °C). Subsequently, cells were washed three times with PBS and permeabilized with 0.5% Triton X-100 (Sigma, Cas. # 9036-19-5) in PBS and incubated at room temperature for 10 minutes. Next, the cells were washed three times with PBS and blocked with 2% Bovine Serum Albumin (BSA) Fraction V (Roche, Lot. # 69453726) in PBS and incubated at room temperature for 60 minutes. The cell supernatants were removed, and cells were directly incubated with primary antibody BCoV-Nucleocapsid mouse anti-bovine monoclonal (Invitrogen, Clone: FIPV3-70; Cat. # MA1-82189) diluted in blocking solution at 1:100 for overnight at 4°C in dark. On the next day, the lysates were removed, and the cell monolayers were washed three times with PBS, followed by incubation with a secondary antibody goat anti-mouse conjugated with Alexa-Flour 488 (Invitrogen, Clone: gG2a (y2a), Ref. # A21131) in BSA blocker at the desired concentration (1:200) for 1 hour at room temperature in the dark. The cells were washed three times with PBS and counterstained with DAPI (Invitrogen, Ref. # D1306) for 5 minutes at room temperature at the desired concentration suggested by the manufacturer in PBS. The cells were washed and examined immediately under fluorescent microscope images at 20x magnification (ZEISS LSM 900).

### 2.6. The Sodium Dodecyl Sulfate-Polyacrylamide Gel Electrophoresis (SDS-PAGE) and the Western blot assay (WB)

The collected proteins from the BCoV-13 and the sham-infected MDBK cells were lysed with the Radio Immuno-Precipitation Assay (RIPA) lysis buffer (Thermo Scientific, Ref. # 89901), supplemented with 1% of 0.5 M EDTA solution and 1% of protease and phosphatase inhibitor (Thermo Scientific). The protein concentrations were measured using the BCA kit (Thermo Scientific) following the manufacturer’s instructions. For the SDS-PAGE assay, equal amounts of the 2x Laemmli buffer 2x laemmli sample buffer (Bio-Rad, Cat. # 1610737) were added into the protein samples and incubated at 100°C for 10 minutes. The protein samples were loaded into the SDS-PAGE gel with iBright Prestained Protein Ladder (Invitrogen, Lot. # 2666356) and run for 90 minutes at 100 volts. The total proteins were detected by using the SDS-PAGE analysis stained with Brilliant Blue R (Sigma, Lot. # MKCJ2486). The proteins were visualized using the GelDoc Go Imaging System (Bio-Rad). The BCoV-13 structural proteins’ molecular weight was determined using an online protein molecular weight calculator (https://www.bioinformatics.org/sms/prot_mw.html). For the Western blot analysis, the protein from the SDS gel was transferred into a Polyvinylidene Difluoride (PVDF) membrane (Bio-Rad, Cat. # 1620177). The PVDF membranes were blocked with 5% BSA diluted in Tris-buffered saline (TBS) containing 0.05% Tween-20 (TBST) and incubated for 1 hour at room temperature. Then, primary antibodies were added into the 5% BSA solution with the desired concentration recommended by the manufacturer and incubated overnight at 4°C. On the next day, the membranes were washed three times with TBST and incubated with the horseradish peroxidase (HRP)-conjugated secondary antibodies diluted in the blocking reagent at the desired concentration recommended by the manufacturer for 1 hour at room temperature. Finally, the membranes were washed with TBST three times, followed by soaking with Clarity Western Enhanced Chemiluminescence (ECL) Substrate (Bio-Rad, Cat. # 170-5060) and visualized using the GelDoc Go Imaging System (Bio-Rad). The primary antibodies used were BCoV-Nucleocapsid (BCoV-N) mouse anti-bovine monoclonal (Clone: FIPV3-70, Cat. # MA1-82189), the BCoV-Spike (BCoV-S) rabbit anti-bovine polyclonal (Cat. # PA5-117562), and the β-actin rabbit anti-bovine polyclonal (Cat. # PA1-46296). The secondary antibodies used were the HRP-conjugated IgG (H+L) for the goat anti-rabbit (Ref. # 31460) or the goat anti-mouse (Ref. # 31430). All the primary and secondary antibodies were purchased from Invitrogen.

### 2.7. The Next-Generation Sequencing (NGS) sample processing, library preparation, and RNA Sequencing techniques

The extracted RNAs from the BCoV-13 were treated with DNase I (RNase-free) (New England BioLabs; Lot. # 10213692) following manufacturer instructions. The RNA sequencing was performed at Azenta Life Sciences (South Plainfield, NJ, USA). In brief, RNA sequencing libraries were constructed with the NEBNext Ultra II RNA Library Preparation Kit for Illumina, following the manufacturer’s recommendations. Sequencing libraries were validated using the Agilent Tapestation 4200 (Agilent Technologies, Palo Alto, CA, USA) and quantified using Qubit 2.0 Fluorometer (ThermoFisher Scientific, Waltham, MA, USA) as well as by quantitative PCR (KAPA Biosystems, Wilmington, MA, USA). Samples were sequenced on a NovaSeq X Plus using a 2x150bp configuration.

### 2.8. Bioinformatic analysis of the NGS data

The raw sequence reads were trimmed to remove possible adapter sequences and nucleotides with poor quality using Trimmomatic v.0.36. The reads were then mapped to the Multiple coronavirus reference genomes available on NCBI using bwa v.0.7.12. The SNPs/INDELs were detected using sam-tools mpileup in conjunction with VarScan v.2.3.9. The settings of VarScan are minimum coverage: 10, minimum reads 4, minimum variant frequency: 0.5%, minimum p-value: 0.05. One file was generated for the variants detected in each sample. The variant files were used with the reference genome sequence (BCoV-Mebus strain) to generate consensus sequences for each sample in .fasta format using the bcf-tools consensus tool v.1.8 standard settings.

### 2.9. GenBank accession numbers

The Genbank accession numbers used in this study were shown in the phylogenetic trees (Figure 6-9).

### 2.10. The Multiple Sequence Alignment (MSA) and Phylogenetic Analysis

The multiple sequence alignment was conducted using the obtained sequences of the BCoV-13 isolate, including (S, N, HE, and the full-length genome) performed using Geneious Prime 2024 Version 11.0.20.1 (https://www.geneious.com) [32]. Our MSA included several representatives per BCoV genotype [25]. The confidence limits of the phylogenetic tree branches were assessed using bootstrap values derived from 1000 replicates. The multiple sequence alignment and phylogenetic analysis were performed based on BCoV isolates from Asia, Europe, and North America from 1972 to 2022, including the BCoV-13 strain isolated from Iowa, USA in 2024 (from this study). All the phylogenetic trees were constructed using MEGA-11 software as previously described [32]. The Human Coronavirus OC43 (HCoV-O43) isolate LRTI 238 from Mexico in 2011 was used as an out-group. The BCoV Gene S, N, HE, and complete BCoV genome reference strains were obtained from the National Center for Biotechnology Information (NCBI) GenBank database. The NCBI accession numbers, country, and year of isolation of reference strains are displayed in Figures 4, 5, 6, and 7.

### 2.11. Statistical analysis

All the data are presented using the mean ± standard deviation (SD). All the statistical analysis was performed using GraphPad Prism Version 9 (http://www.graphpad.com/faq/viewfaq.cfm?faq=1362, released on Oct 27, 2020). The paired comparison was performed using Student’s t-test as previously described [33]. The statistical significance was set at P values less than 0.05. The statistical significance is denoted as: (* P < 0.05, ** P < 0.01, *** P < 0.001, and **** P < 0.0001).

## 3. Results

### 3.1. Propagation and isolation of the BCoV field isolates

The BCoV-13 isolate infection in Human Rectal Tumor-18 (HRT-18) cells triggered an apparent cytopathic effect (CPE) and morphological changes, including cell rounding and detachment from the monolayer at 72 hours post-infection (hpi) compared to the sham (Figure 1A, 1B). The CPE in the MDBK cells was noticeable after 96 hpi (Figure 1D, 1E). Morphological examination revealed that the CPE effect was more prominent in BCoV-13 passage 5 (P-5) compared to the P-2 in HRT cells (Figure 1B, 1C). Similarly, MDBK cells infected with the BCoV-13 P-5 displayed a more prominent CPE compared to those infected with cell culture supernatants from BCoV-13 P-2 (Figure 1E, 1F).

**Figure 1:**
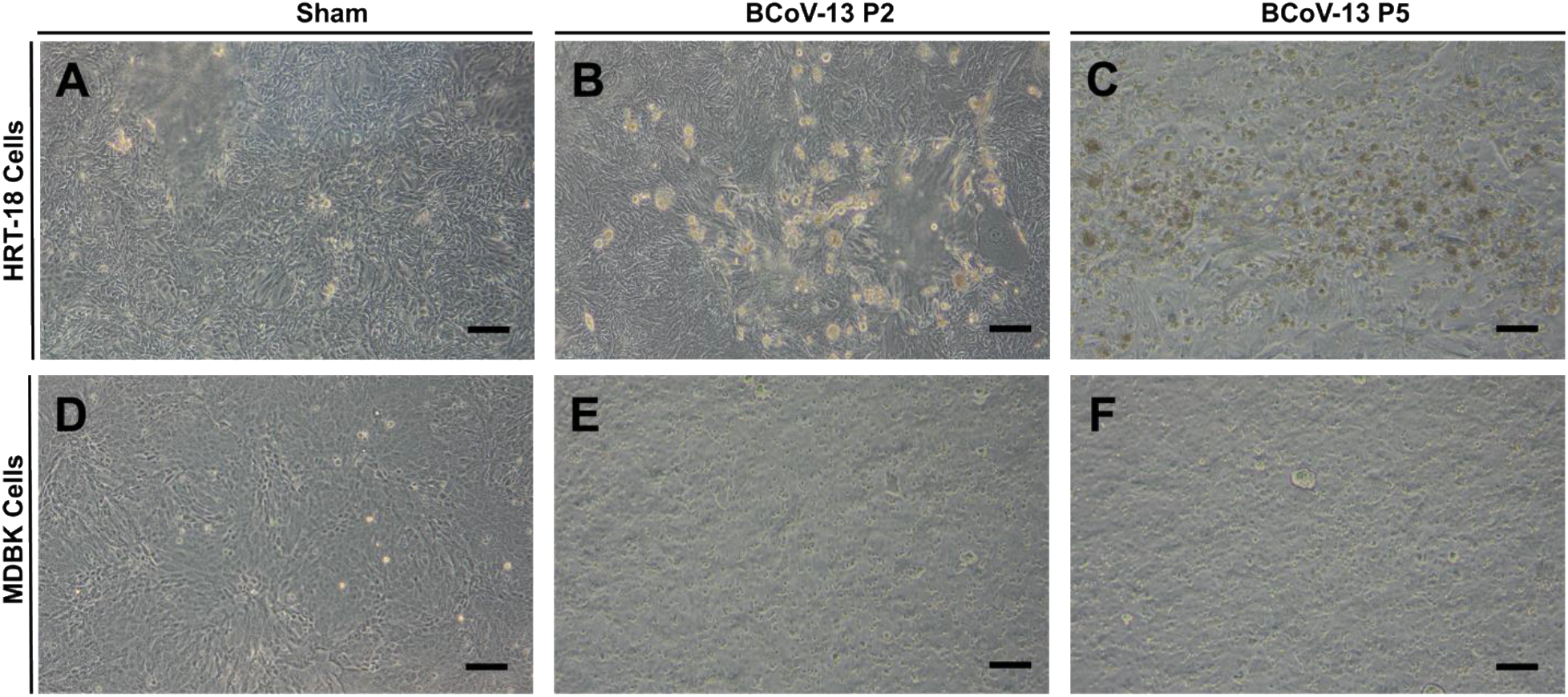
(A) Morphological observation of HRT-18 cell line in Sham (control), and (B) cytopathic effect (CPE) at 72 hours post-infection (hpi) in BCoV-13 P-2 infected HRT cells. (C) CPE at 72 hpi in BCoV-13 P-5 infected HRT cells. (D) Morphological observation of MDBK cells in Sham, (E) and CPE at 96 hpi in BCoV-13 P-2 infected MDBK cells. (F) CPE at 96 hpi in BCoV-13 P-5 infected MDBK cells.

### 3.2. Confirmation of BCoV-13 infection and replication *in-vitro* by the IFA

BCoV-13 infection in the MDBK cells was observed in several subsequent passages by IFA using the BCoV-N fluorescent conjugated antibody. The BCoV infection was identified by the green fluorescent signal in the infected cells (Figure 2B, 2E). The IFA results demonstrated a higher BCoV-N protein expression in P-5 compared to the P-2 (Figure 2B, 2C, 2E, 2F), suggesting a significant increase in the viral production with subsequent blind *in-vitro* passages of this viral isolate.

**Figure 2:**
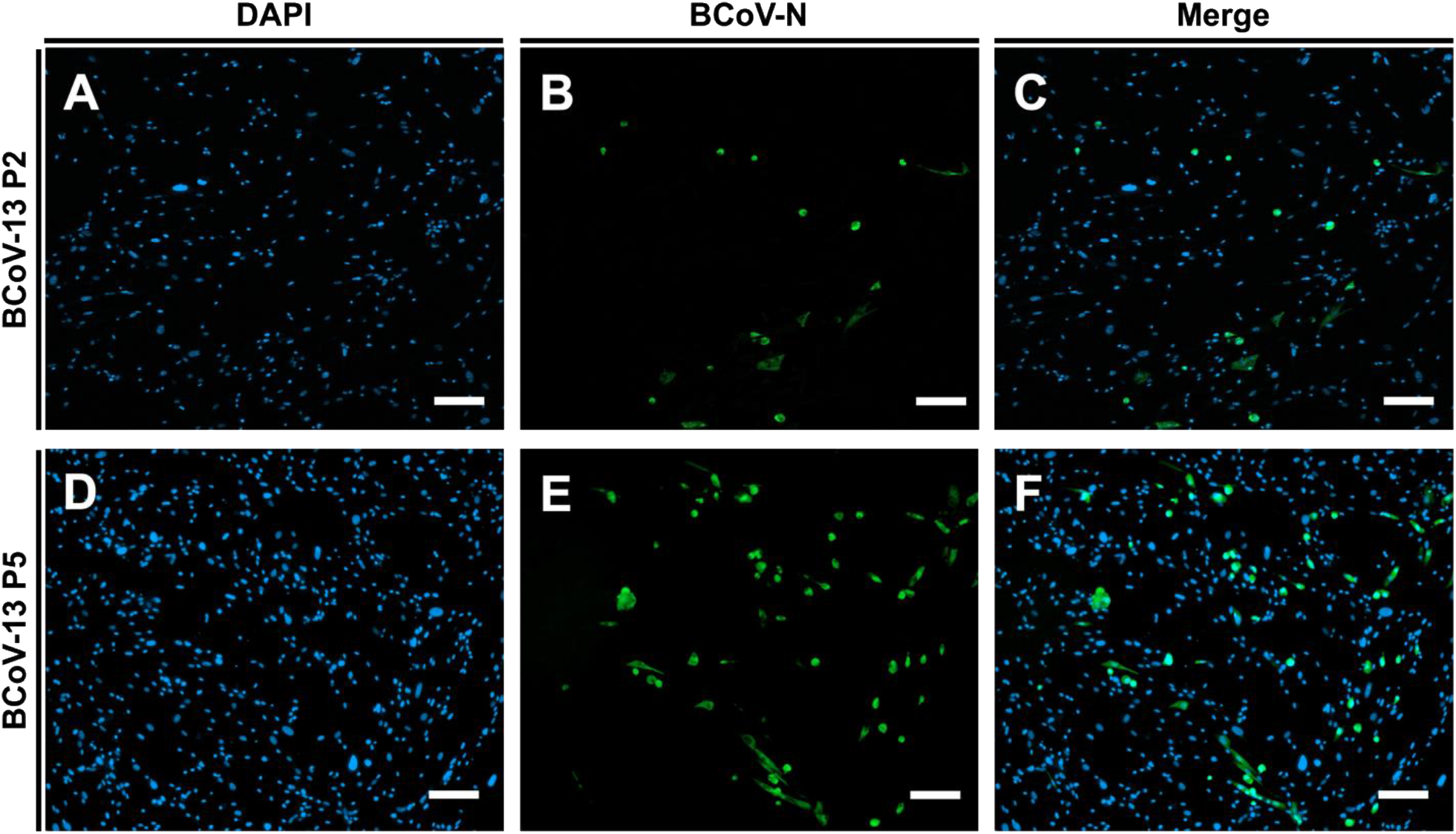
Immunofluorescence (IF) staining of the MDBK cells infected with BCoV-13 P-2; (A) DAPI (blue color), (B) BCoV-Nucelocapsid antibody (green color), and (C) Merge. IFA results of the MDBK cells infeted with BCoV- 13 P-5; (D) DAPI (blue color), (E) BCoV-Nucelocapsid antibody (green color), and (F) Merge. MDBK cells were infected with BCoV-13 at different passages, and IF analysis was performed after 72 hpi.

### 3.3. Quantification of the BCoV-13 genome copy numbers and the viral infectivity titers in different passages in the HRT and the MDBK cells

The qRT-PCR was performed on the original BCoV-13 sample collected from the fecal sample, and the presence of BCoV was confirmed (Figure 3A). The BCoV-13 genomic copy numbers in P-5 were significantly higher compared to P-2 in the MDBK and the HRT-18 cell lines (Figure 3B). Similarly, the BCoV-13 viral infectivity titers showed approximately 138.5 times higher in cell culture supernatants collected from P-5 than that of P-3 in both MDBK and HRT-18 cells (Figure 3C). The average plaque size of BCoV-13 P-2 was measured to be 0.39 mm in diameter, while the average plaque size of BCoV-13 P-5 was 0.53 mm (Figure 3D). The plaque size of BCoV-13 P-5 was significantly larger than BCoV-13 P-2.

**Figure 3:**
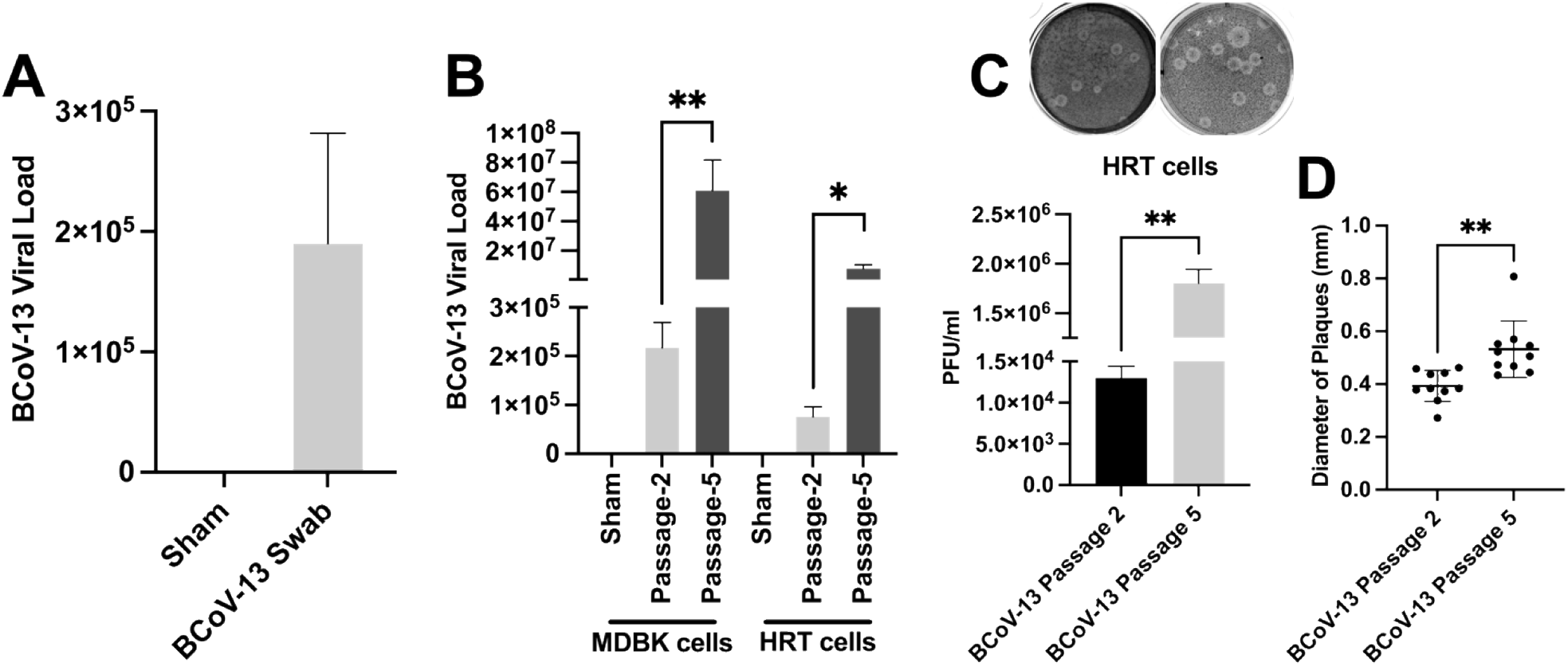
Quantitative estimation of the BCoV-13 genome copy number and infectivity titers by the qRt-PCR and the viral plaque assay, respectively. (A) The qRT-PCR analysis shows the genome copy numbers of the original sample of the BCoV-13 isolate. (B) The qRT-PCR analysis of the BCoV-13 P2 and P5 genome copy numbers in MDBK and HRT cell lines. (C) Results of the viral plaque assay of cell culture supernatants infected with P2 and P5 in the HRT- 18 cell lines. (D) The plaque diameter of BCoV-13 infected with P2 and P5 was measured on 10 plaques per virus.

### 3.4. Detection and confirmation of the major structural proteins of the BCoV-13 isolate

The SDS-PAGE results of the expressed proteins in the sham and the BCoV-13 infected cells. Results show the protein bands of the BCoV-13 isolate as follows: Spike subunit 1 (S1; ∼84 kDa), Spike subunit 2 (S2; ∼65 kDa), Nucleocapsid (BCoV-N) protein (∼49 kDa), Hemmaglutinin-Esterase (HE) protein (∼47 kDa), Membrane (M) protein (∼26 kDa) and Internal (I) protein (∼22 kDa) (Figure 4A). Confirmation of the expression of the BCoV-N and the BCoV-S glycoprotein was confirmed by the western blot analysis (Figure 4B).

**Figure 4:**
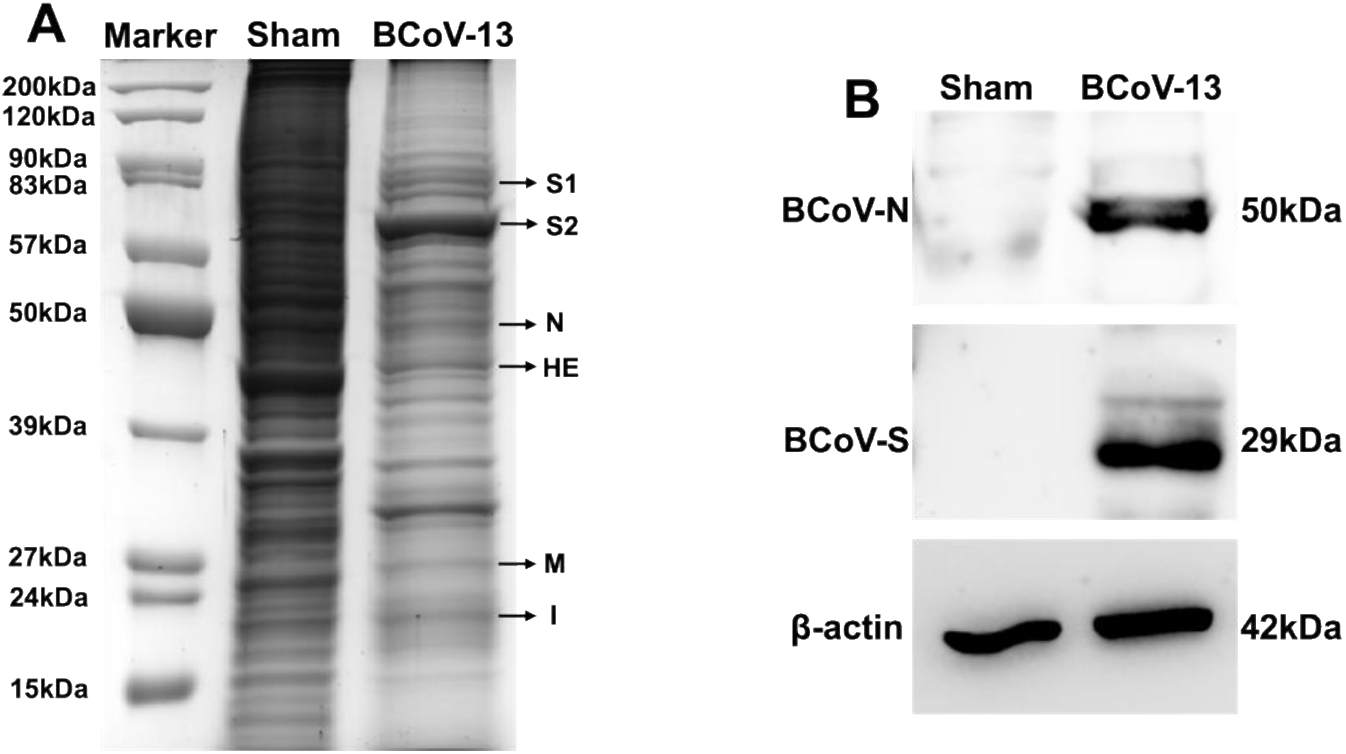
(A) Sodium Dodecyl Sulfate-Polyacrylamide Gel Electrophoresis (SDS-PAGE) results showing the total protein in sham and BCoV-13 infected MDBK cells. Lane-1 is the protein marker; Lane-2 is sham MDBK cells; Lane-3 is MDBK cells infected with BCoV-13. **(B)** Western blot analysis of BCoV-nucleocapsid (BCoV-N) protein, BCoV-spike (BCoV-S) protein, and β-actin protein expression in MDBK cells infected with sham and the BCoV-13 infected cells.

### 3.5. Establishment of the genome structure and organization of the BCoV-13 isolate

The full-length genome sequence of the BCoV-13 isolate was drafted as described above through the NGS and deposited under the GenBank (accession no # **PQ118397**). The viral genome size of this isolate is 31022 nucleotides. The GC contents of the genome of the BCoV-13 isolate is (37%). The viral genome is flanked with two untranslated regions at their 5’ and 3; ends. The 5’ two-thirds of the genome is occupied with a large gene called Gene-1, which consists of two large overlapping open reading frames (ORFs) with a ribosomal frameshifting mapped at position 13341 (Figure 5A). The ORF1a/b is the longest BCoV gene of 21284 nucleotides composed of 16 Non-structural proteins (NSPs) (Figure 5B). ORF1a contains 11 NSPs, and ORF1b contains 5 NSPs. NSP 11 and 12 overlap ORF1a and ORF1b (Figure 5B). The size, start, and stop position of NSPs are shown in Figure 5B. The 3’ one-third of the genome contains five main structural proteins (HE. S, M, E, and N) interspersed with some non-structural proteins (32kDa, 4.9kDa, 4.8kDa, and 12.7kDa) (Figure 5A). Meanwhile, a small internal (I) protein of 615 base pairs (bp) is mapped inside the N protein (Figure 5A). The Spike glycoprotein (S) gene is 4092 nucleotides in length, composed of subunits the S1 and the S2 subunits (Figure 5C). The full-length S protein is cleaved by the Furin host serine protease; the cleavage site is mapped between the S1 and S2 subunits at position 25930 (Figure 5C). The S1 subunit contains the Receptor Binding Domain (RBD), N-terminal domain (NTD), and C-terminal domain (CTD) (Figure 5C). The S2 subunit is composed of the Fusion Peptide 1 (FP1), Fusion Peptide 2 (FP2), Heptad Repeat 1 (HR1), and Heptad Repeat 2 (HR2) (Figure 5C). The Nucleocapsid (N) gene of BCoV-13 is 1338 nucleotides in length (Figure 5D). The N gene comprises the RNA-Binding Domain, NTD, CTD, and Dimerization Domain (Figure 5D). A notable deletion of 9 bp was found in BCoV-13 at the linker position 620, immediately downstream of the RNA-Binding Domain (Figure 5D).

**Figure 5:**
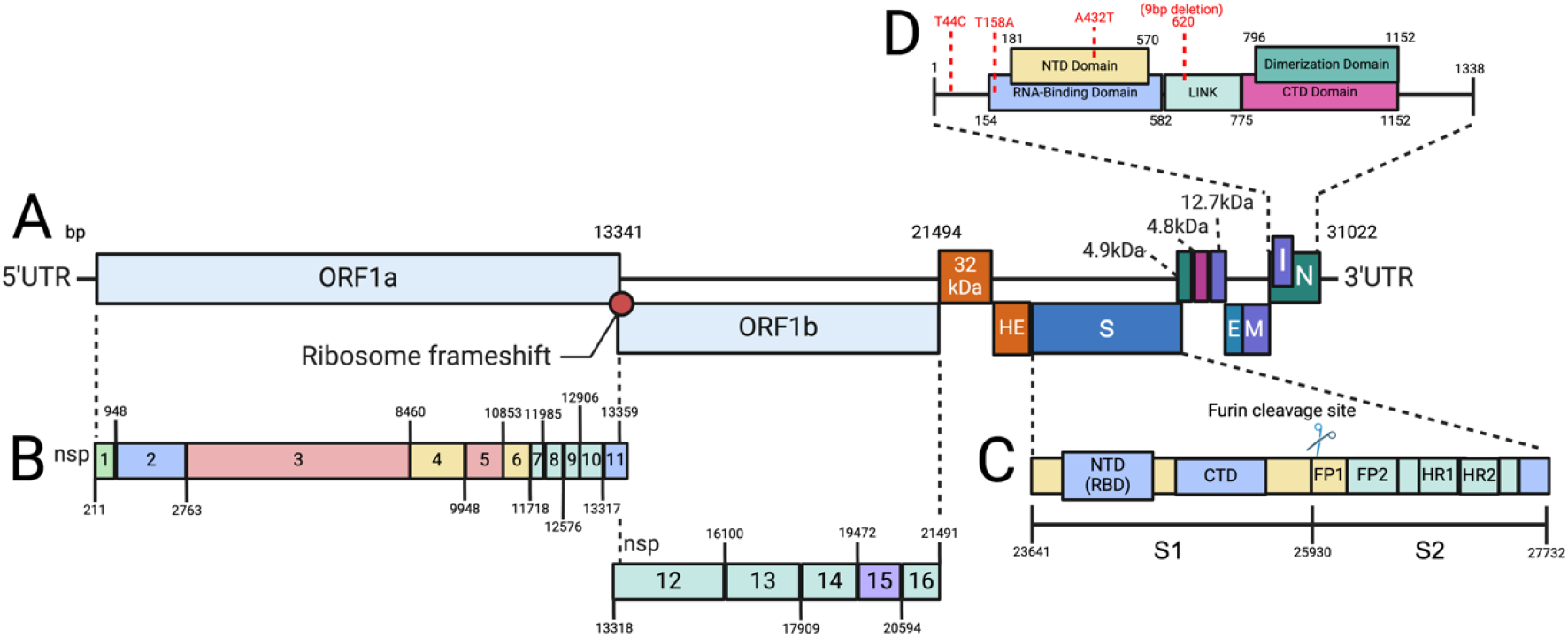
The BCoV-13 genome structure and organization. (A) A schematic representation of the genome structure and organization of the BCoV-13 isolates based on full-length genome sequencing. The genome is composed of 5’Un-Translated Region (UTR), Open Reading Frame (ORF)-1a, ORF1b, 32kDa, Hemmaglutinin- Esterase (HE), Spike (S), 4.9kDa, 4.8kDA, 12.7kDa, Envelope (E), Membrane (M), Nucleocapsid (N), Internal (I), and 3’UTR. **(B)** A schematic representation of the 16 Non-Structural Proteins (NSP) within the ORF1a/b showing their relative sizes and positions. **(C)** A schematic representation of the Spike glycoprotein (S) showing their relative subunits, sizes, and positions. S1 subunit and S2 subunit of spike gene, with a Furin cleavage site between S1 and S2 subunit. S1 contains Receptor Binding Domain (RBD), N-terminal domain (NTD), and C-terminal domain (CTD), and the S2 subunit contains Fusion Peptide 1 (FP1), Fusion Peptide 2 (FP2), Heptad Repeat 1 (HR1), and Heptad Repeat 2 (HR2). **(D)** A schematic representation of the Nucleocapsid (N) gene showing their relative domains, sizes, and positions. The N gene is composed of RNA-Binding Domain, NTD, CTD, and Dimerization Domain.

### 3.6. A comparative analysis of the BCoV-13 genome sequencing with the reference BCoV Enteric and Respiratory isolates

The BCoV-13 genome shows 99.8% similarity to the BCoV enteric Mebus isolates and 98.8% with the BCoV-LUN respiratory isolate. A detailed comparative analysis identified nucleotide variations between BCoV-13, Mebus, and BCoV-LUN isolates (Table 1). The results revealed 27 nucleotide variations between BCoV-13 and the Mebus isolate. Among these variations, 30 nucleotide changes were observed in ORF1ab and 12 variations in the spike gene. Only one nucleotide variation (G22658C) was reported in the HE gene. In the nucleocapsid gene, variations were observed at four different sites, including a notable deletion of 9 base pairs (bp) at position 30014 in BCoV-13 (Table 1). One nucleotide deletion was also observed in the 3’UTR region, resulting in a genome size of 31022 bp for the BCoV-13.

**Table 1:**
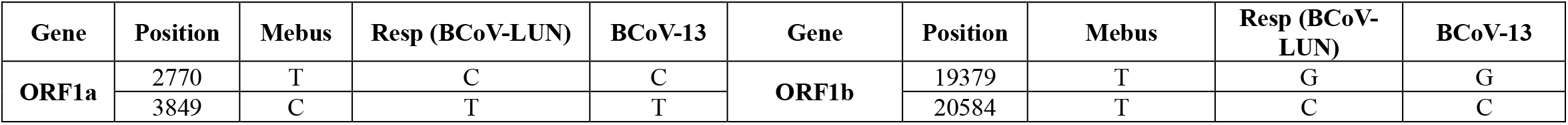
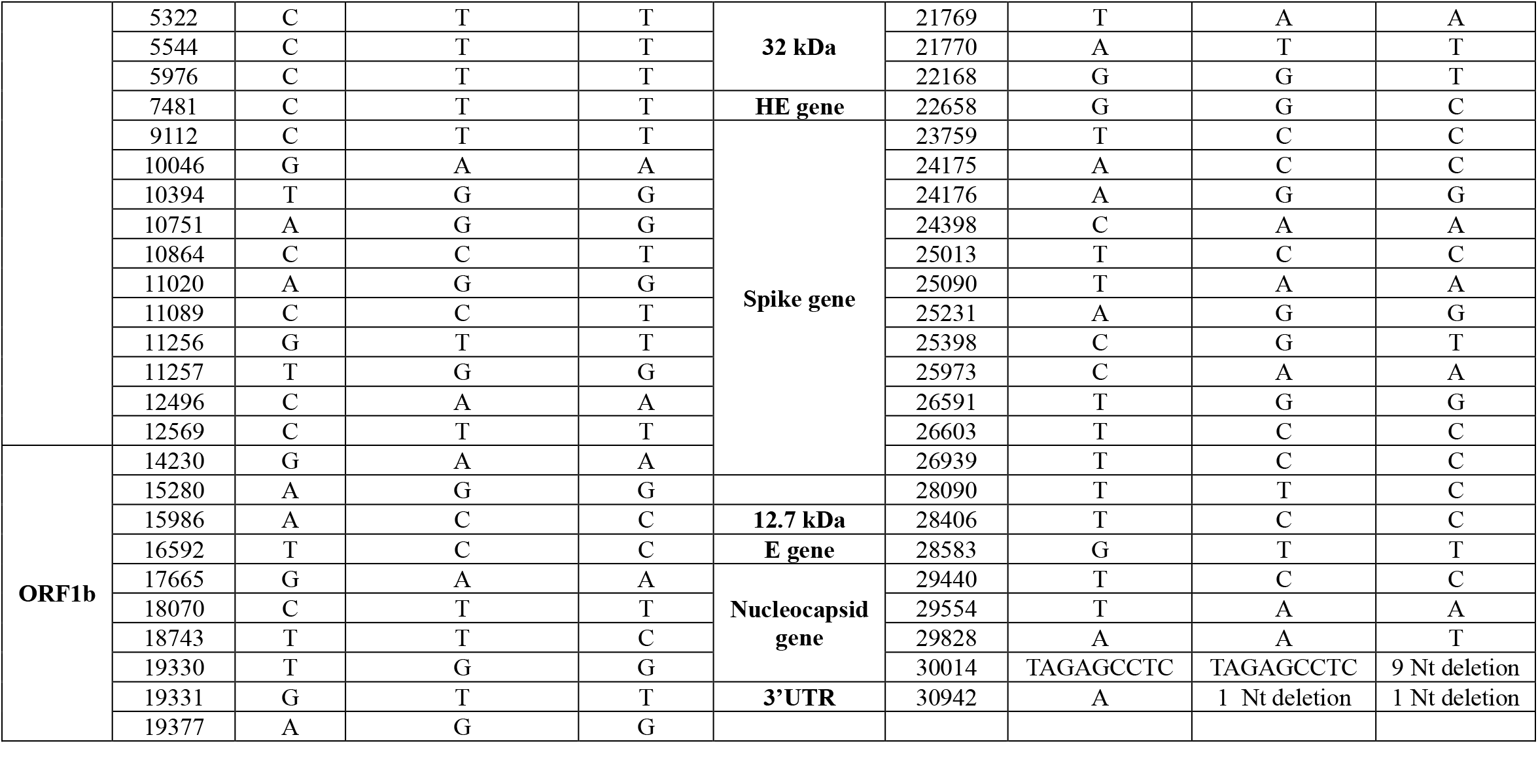
Mapping the variations of the nucleotide sequences among the full-length genome of the BCoV-13 isolate compared to the Mebus and BCoV-LUN isolates.

### 3.7. The phylogenetic analysis of the BCoV-13 based on the nucleotide sequences of the nucleocapsid protein gene

The phylogenetic tree was based on the nucleotide sequences of the nucleocapsid gene, and the tree was split into two groups (GI and GII). The GI group includes the old classical isolates from Japan, Canada, the USA, India, and Taiwan. The BCoV-13 isolate reported in this study clustered with the sequences of the group GI together with other classical BCoV isolates (Figure 4). The GII was further divided into two subgroups: GIIa (North America and Asia) and GIIb (Europe). The group GIIa includes BCoV isolates from the USA between 1998 and 2022, including some classical isolates like BCoV-Mebus and BCoV-LUN (Figure 6). Group GIIa also includes recent BCoV isolates from China and Japan reported between 2015 and 2022 (Figure 6). The group GIIb includes BCoV isolates reported from Europe, including France, Italy, Ireland, and Tukey, between 2013 and 2019 (Figure 6).

**Figure 6:**
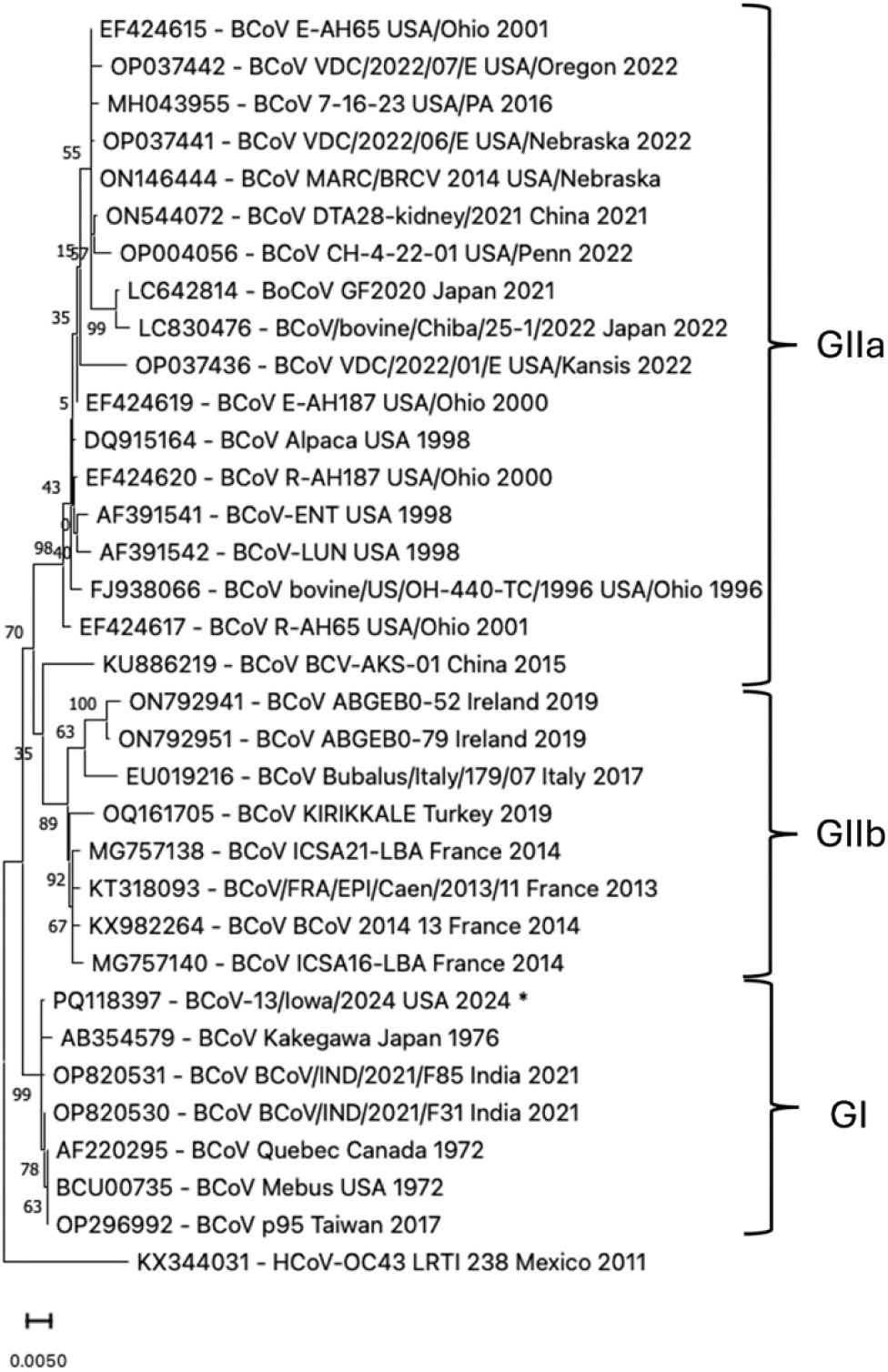
The Phylogenetic tree based on the Nucleocapsid gene sequences from 34 BCoV isolated worldwide, including the BCoV-13 isolate detected in this study, shown with an asterisk (*). The Phylogenetic tree with the highest Log Likelihood (-3079.82) was constructed using the maximum likelihood method and the Tamura-Nei model with bootstrap values (1000 replicates) using MEGA 11 software. The percentage of the tree in which the associated taxa clustered together is shown next to the branches. The genotype distribution of the isolates is shown on the right side of the figure. The Human Coronavirus (HCoV-OC43) was used as an out-group control.

### 3.8. The phylogenetic analysis of the BCoV-13 based on the nucleotide sequences of the Spike glycoprotein

The phylogenetic analysis revealed that spike glycoprotein of BCoV-13 isolate was included in group GI (Figure 7). In addition, the GI group consists of the classical BCoV Isolate Kakogawa, Japan 1976, BCoV Quebec Canada 1972, and BCoV Mebus USA 1972 (Figure 7). The GI group also includes a BCoV V270 from Germany in 2006 and a recent isolate from Taiwan and India in 2017 and 2021, respectively (Figure 7). The group GIIa contains BCoV isolates from the USA, China, Japan, and Korea between 1996 and 2022 (Figure 7). The group GIIb includes BCoV isolates from Europe, including France, Italy, Ireland, Sweden, and Tukey between 2005 and 2019 (Figure 7).

**Figure 7:**
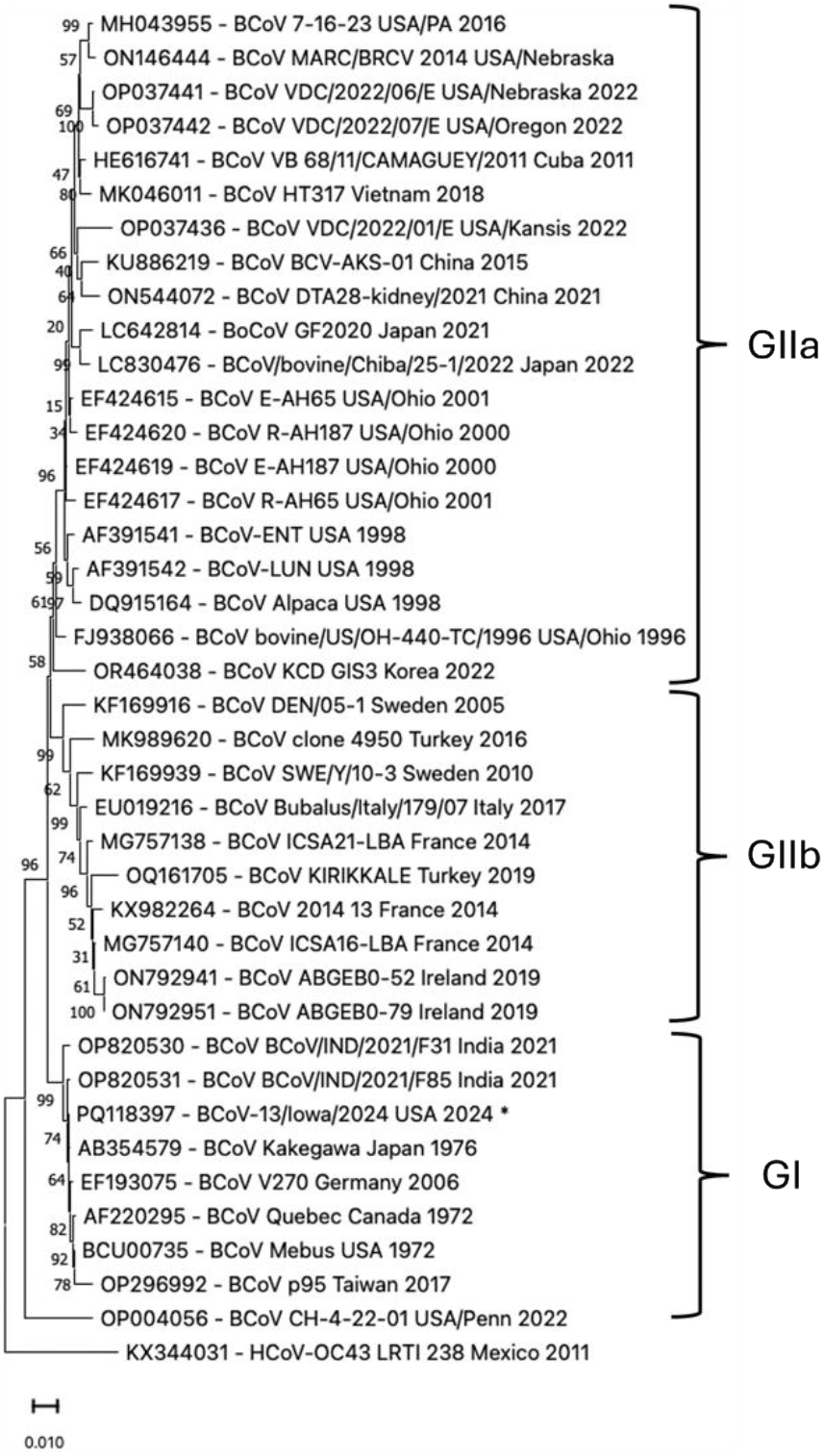
The phylogenetic tree based on the spike glycoprotein gene sequences from 40 BCoV isolates worldwide, including the BCoV-13 isolate detected in this study, shown in steric. The Phylogenetic tree with the highest Log Likelihood (-13605.05) was constructed using the maximum likelihood method and Tamura-Nei model with bootstrap values (1000 replicates) in MEGA 11 software. The percentage of the trees in which the associated taxa clustered together is shown next to the branches. The genotype distribution of the isolates is shown on the right side of the figure. The Human Coronavirus (HCoV-OC43) was used as an out-group control.

### 3.9. The Phylogenetic analysis of the BCoV-13 based on the nucleotide sequences of the HE gene

The phylogenetic analysis based on the nucleotide sequences of the HE gene revealed that the BCoV-13 isolate was clustered within the GI along with the classical BCoV Isolate from Quebec, Canada 1972, BCoV Mebus USA 1972, BCoV Kakogawa Japan 1976 (Figure 8). The GI group also includes the BCoV-A3 and the SUN5 isolate from Korea reported in 1994, and a recent isolate from Taiwan and India in 2017 and 2021, respectively (Figure 8). The group GIIa contains BCoV isolates from the USA, China, Vietnam, Japan, and Korea between 1998 and 2022 (Figure 8). The group GIIb includes BCoV isolates from Europe, including France, Italy, Ireland, and Tukey, between 2014 and 2019 (Figure 8).

**Figure 8:**
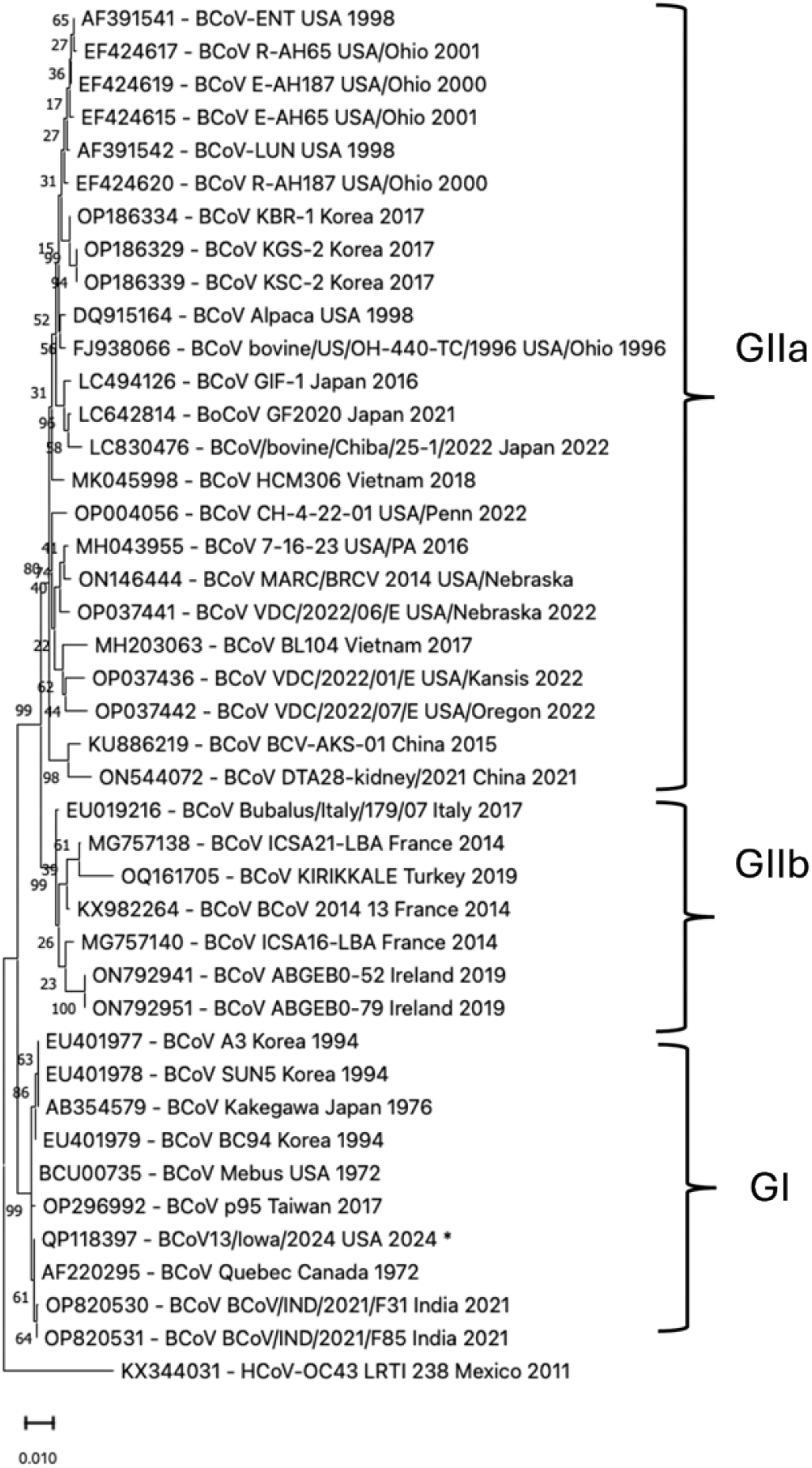
The Phylogenetic tree based on the HE gene sequences from 42 BCoV isolates worldwide, including the BCoV-13 isolate detected in this study shown in steric. The Phylogenetic tree with the highest Log Likelihood (- 3456.64) was constructed using the maximum likelihood method and Tamura-Nei model with bootstrap values (1000 replicates) in MEGA 11 software. The percentage of the trees in which the associated taxa clustered together is shown next to the branches. The genotype distribution of the isolates is shown on the right side of the figure. The Human Coronavirus (HCoV-OC43) was used as an out-group control.

### 3.10. The Phylogenetic analysis of the BCoV-13 based on the nucleotide sequences of the full-length genome

The phylogenetic analysis of the full-length genome of BCoV isolates revealed that the BCoV-13 isolate was clustered with the other BCoV isolates in group GI, including some classical BCoV isolates from Quebec, Canada 1972, BCoV Mebus USA 1972, and BCoV Kakogawa Japan 1976 (Figure 9). The GI group also includes two isolates, F31 and F85, from India in 2021 and one isolate from Taiwan p95 in 2017 (Figure 9). The group GIIa contains BCoV isolates from the USA, China, and Japan between 1996 and 2022 (Figure 9). The group GIIb includes BCoV isolates from Europe, including France, Ireland, and Tukey, between 2014 and 2019 (Figure 9).

**Figure 9:**
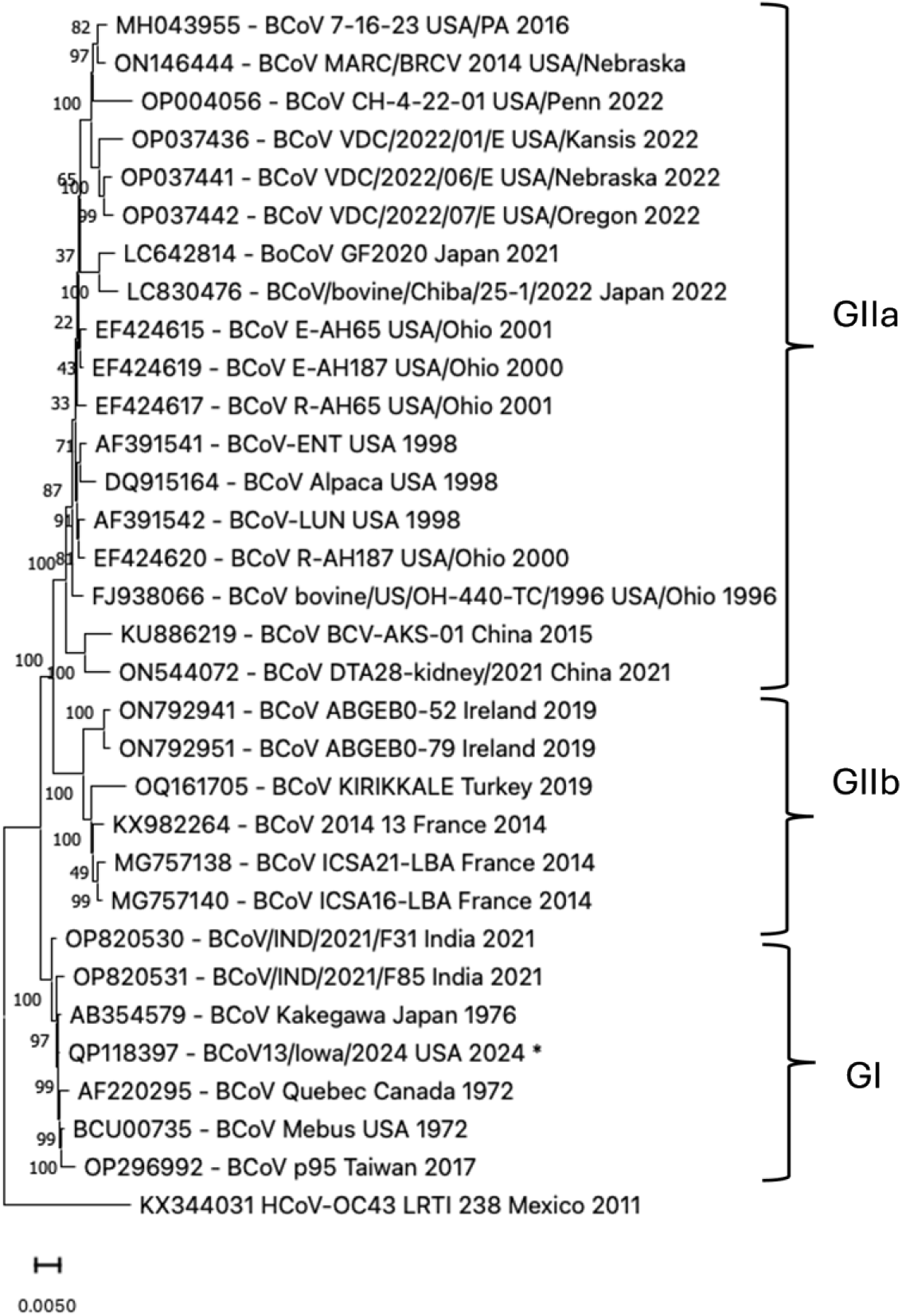
The Phylogenetic tree based on the nucleotide sequences of the BCoV-13 isolate full-length genomes from 32 BCoV isolates worldwide, including the BCoV-13 isolate detected in this study shown in steric. The Phylogenetic tree with the highest Log Likelihood (-74227.53) was constructed using the maximum likelihood method and Tamura-Nei model with bootstrap values (1000 replicates) in MEGA 11 software. The percentage of the tree in which the associated taxa clustered together is shown next to the branches. The genotype distribution of the isolates is shown on the right side of the figure. The Human Coronavirus (HCoV-OC43) was used as an out-group control.

### 3.11. Mapping the mutations across the BCoV-13 spike glycoprotein

Based on the pairwise sequence alignment of the BCoV-13 isolate with the BCoV Mebus isolate, the spike glycoprotein gene of BCoV-13 showed 12 nucleotide mutations (Table 1 and Figure 10). Four nucleotide mutations were observed in the N-terminal domain (NTD) of the S1 Subunit (Figure 10). The first substitution at the NTD **T**23759**C** changed the amino acid *Isoleucine* to *Threonine* (**I** 40 **T**). Another substitution of two consecutive nucleotides at A24175C and A24176G was observed at the NTD region, changing the Lysine into Arginine (K179R). The fourth nucleotide substitution is located at the NTD C24398A, leading to the amino acid Serine change into Tyrosine (S253Y) (Figure 10). Four nucleotide mutations were observed at the C-terminal domain (CTD) of the S1 subunit. The substitution T25013C at the CTD changed the amino acid Phenylalanine into Serine (F458S). The T25090A substitution at the CTD changed the amino acid Serine into Threonine (S484T). The A25231G substitution at the CTD changed the Asparagine into the Aspartic acid (N531D). The fourth substitution was the C25398T at the CTD, which showed no change at the amino acid level. Four nucleotide substitutions were observed at the S2 subunit of the spike glycoprotein gene (Figure 10).

**Figure 10:**
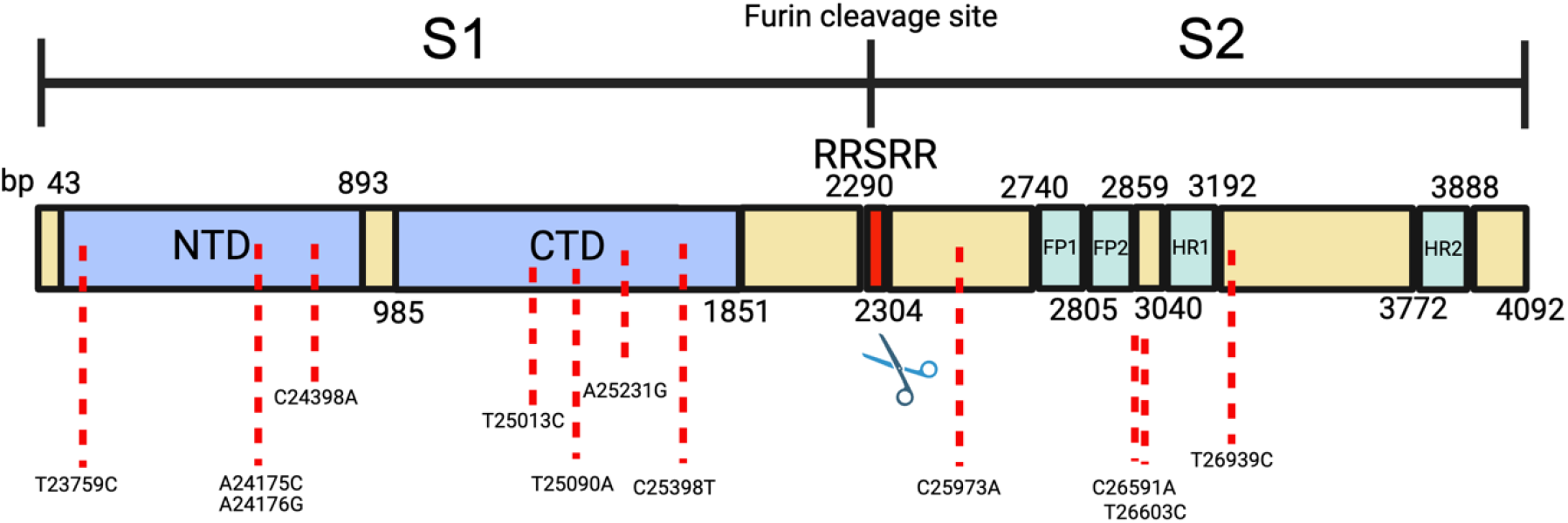
A schematic representation of the structure and organization of the BCoV-13 isolate spike glycoprotein and mapping of the mutations among various domains. The structure of the (S) gene shows their relative subunits, their sizes, and mapping their locations. The S1 and S2 subunits of the spike gene show the Furin cleavage site (RRSRR amino acids) separating the S1 and S2 subunits. The S1 contains the Receptor Binding Domain (RBD), the N-terminal domain (NTD), and the C-terminal domain (CTD). The S2 subunit contains the Fusion Peptide 1 (FP1), Fusion Peptide 2 (FP2), the Heptad Repeat 1 (HR1), and the Heptad Repeat 2 (HR2). The nucleotide substation is shown with a red dotted line in each section. The scale of the spike gene is shown in base pair (bp) at each position.

### 3.12. Notable deletion in the BCoV-13 isolate downstream the RNA binding domain of the nucleocapsid protein

The Nucleocapsid gene of BCoV-13 showed three nucleotide mutations and nine-nucleotide deletion based on the pairwise alignment with the Mebus isolate (Table 1). The first substitution, T29440C, was observed in the downstream RNA-binding domain (RBD) of the N gene, which resulted in the change of the Phenylalanine into Serine (F15S) (Figure 5D). The T29554A substitution was observed in the RNA-Binding domain (RBD) of the N gene, changing the Leucine into Glutamine (L53Q) (Figure 5D)—another substitution, A29828T, at the RBD of the N gene. The multiple sequence alignment of 33 different BCoV isolates and one HCoV-OC43 from various countries revealed that nine nucleotide deletions were found exclusively in BCoV-13 (Figure 11). These nine nucleotide deletions showed three amino acid deletions 207-Arginine-Alanine-Serine- 209 (207RAS209) in BCoV-13 compared with Mebus.

**Figure 11:**
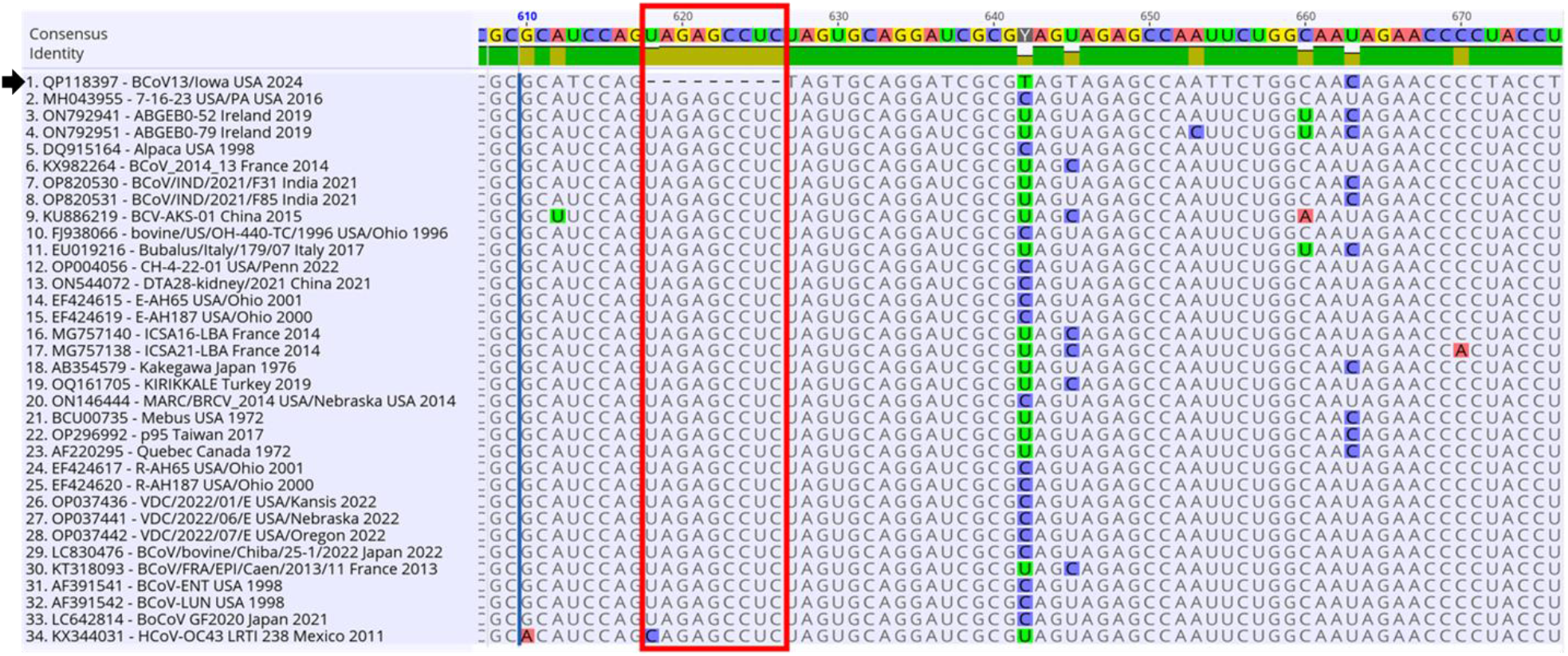
The Multiple Sequence Alignment (MSA) of Nucleocapsid gene sequences from 33 BCoV strains and one HCoV-OC43, isolated worldwide. The black arrow indicates the BCoV-13 isolate reported in this study. The 9- nucleotide deletion region is highlighted in the red box. The MSA was performed using Geneious Prime 2024 Version 11 software.

## 4. Discussion

BCoV infection continues to negatively impact the cattle industry since its discovery [1, 34, 35]. Despite the availability of some BCoV vaccines, there is a continuous report of the many outbreaks of viral infection among cattle populations across the globe, including the USA [36, 37]. As with other coronaviruses, the BCoV genome is always under dynamic changes because of the mutation frequency due to the lack of the proofreading capability of the RNA polymerase enzyme and the possibility of recombination between different isolates [36, 38]. The S, HE, and N proteins of the BCoV play essential roles in viral infection, replication, and pathogenesis [39]. The BCoV-S protein is a complex protein consisting of three identical units combined to form a homotrimer. BCoV-S is a highly glycosylated protein that helps the virus evade the host immune response. BCoV-S protein is usually cleaved by host cell processes to produce two subunits (S1 and S2) [40, 41]. This cleavage step is crucial in activating the BCoV-S protein to initiate the viral infection. The S1 subunit contains the receptor binding domain (RBD), which is responsible for the attachment of the virus to the host cell. The S2 subunit mediates the fusion between the viral envelope and the host cell membrane to facilitate the entry of the viral genome into the host cell. The BCoV-HE is a homodimer. Its unit comprises the two domains (lectin and the esterase) [42]. The first is responsible for binding the viral receptors, the N-acetyl-9-O-acetylneuraminic. However, the esterase domain cleaves the acetyl group from the 9-O-acetylated sialic acid, facilitating the viral release from the infected cells. The BCoV-HE plays an essential role in viral immune evasion by masking the S protein to avoid its recognition by neutralizing antibodies [43]. The BCoV-N is one of the most conserved proteins in the BCoV. It consists of the (N) terminal domain (NTD) and the (CTD) terminal domain connected with a linker sequence. The NTD is essential for binding the viral nucleic acid, while the CTD assists in the viral packaging [44, 45]. Genotyping of BCoV isolate based on the phylogenetic analysis of the viral sequences, especially using the BCoV-(ORF1a, HE, S, and N) sequences, identified 14 known genotypes [25]. Eleven out of them represent the European isolates, and three represent the North American isolates [25]. A recent study restructured this classification into three main genotypes (GI-GIII) based on the available BCoV genome sequences collected between 987 and 2017 reported from various countries around the globe [46]. The GI is further divided into nine subgroups. The GI-I is the most notable genotype circulating in US cattle [46]. Initially, we tried propagating three BCoV field isolates using the MDBK and HRT-18 cell lines. After three subsequent passages, we focused on only one isolate based on the observed CPEs in the infected cells, the viral genome copy numbers, and the virus infectivity titration by plaque assay. Our results show an increase in the BCoV-13 replication with the progression of the subsequent passages (P2-P5) in cell cultures (Figures 1-3). This observation supports the rise in severities of the observed CPEs in the MDBK and HRT-18 cell lines, the viral genome copy numbers, and the virus infectivity titers by plaque assay (Figures 1 and 3). Also, the BCoV-N protein expression level increased dramatically from P-2 to P5, as shown by the IFA in (Figure 2). It has been demonstrated that the subsequent blind passages of some coronaviruses impacted the virus kinetics and replication. The repeated passages of the virus may result in the adaptation of the virus into this new host and shorten the time required to complete its replication cycle. This frequent passage may also increase the virus infectivity as measured by the plaque assay. The reasons behind this could be the accumulation of several mutations and the production of some viral quasispecies, which lead to more virus particles that lead to intensive CPE in the inoculated cells. Previous studies showed the limited replication potential of some BCoV field isolates to replicate in various cell cultures until several blind passages [47]. The BCoV-13 isolate grew very well in the MDBK and the HRT-18 cell lines with and without the addition of trypsin compared to several filed isolates propagated under the same conditions. On the other hand, the repeated passage of some field isolates in cell culture may attenuate virus infectivity. The best example is the porcine epidemic diarrhea virus (PEDV) attenuation through subsequent passages in the PC22A cell line [48]. However, this attenuation of the PEDV infectivity was achieved after 106 blind passages in the cell line [48].

We believe the BCoV-13 isolate might considered a virulent isolate of the enteric BCoV for several reasons. First, the affected animal died three days after the onset of the clinal signs. Second, this isolate can propagate well in cell culture in the presence or absence of the external source of trypsin. Third, high genome copy numbers, protein expression, and virus infectivity titers were achieved after very few numbers of the passages in cell culture. Fourth, the plaque morphology of the cell culture supernatants collected from P-5 showed a considerable size compared to that of the P-2. Some of the plaques shown in the P-5 fused together to form very large plaques (Figure 3C). Further studies are required to reveal the exact mechanism of the potential BCoV-13 virulence. Despite the reported mutation across the viral genome of the BCoV-13 isolates, our phylogenetic analysis confirms it is still clustered with the other members of the genotype I of the BCoV circulating in North America [4–6, 8, 25]. Our results show several mutations across the BCoV-13 spike glycoprotein, particularly within the RBD (Figure 10). Recent reports showed a similar pattern of mutations in the SARS-CoV-2-RBD, suggesting the plasticity of the viral RBD [49]. Other studies showed that the mutation within the SARS-CoV-2 S1 region, the D614G, increased the viral infectivity and virulence [50]. Other studies showed that some amino acid substitutions within the BCoV-S glycoprotein mapped at locations (146, 148, and 509) resulted in conformational structure changes in the S protein [37]. other studies suggested the amino acids at locations 113, 115, 146, 148, 501, 531, and 646 on the BCoV-S protein could play an important role in the tissue tropism of the virus and may be able to distinguish between the enteric and respiratory isolates of BCoV as novel biological markers [3, 14, 51–54]. In this study, we identified the substitution of Asparagine into Aspartic acid at 531 at the S gene of BCoV-13. Consistent with our results, some BCoV isolates from Turkey showed several mutations across the viral genome, especially F458S, S484T, and N531D mutations [55]. Another study reported a deletion of six amino acids within the BCoV-S protein of some Brazilian isolates at the locations (526-531), which were identical to deletions reported in the HCoV-OC43 [56]. These findings indicate that the mutations in the BCoV-S protein, specifically in the CTD region, could result in some structural changes within the S glycoprotein and alter the viral virulence and pathogenicity in the affected cattle. A recent study showed the emergence of new BCoV isolates with a notable 12 nucleotide insertion in the RBD of the HE gene of some BCoV isolates in the US, suggesting the possibility of further interspecies transmission [49]. This is in contrast to the BCoV-13 HE sequences, which shows only one mutation at the genomic location 22658 (Table 1). These observations suggest the diverse genetic background of the currently circulating strains of BCoV in US cattle. The nucleocapsid protein of coronaviruses plays essential roles in virus replication and immune regulation/evasion and could act as a target for the development of diagnostic assays, antiviral therapy, and vaccines [57–63]. A recent study showed the roles of the BCoV-N protein in the inhibition of the host IFN-β production through the inhibition of the RIG-I-like receptors pathway [64]. Our results show nine nucleotide deletions downstream of the RNA binding domain of the BCoV-13-N protein (Figure 11). This mutation could contribute to the viral immune evasion and the virulence of their BCoV isolate.

However, more functional studies are required to study the roles of these deletions and other mutations across the viral genome for this recent isolate of BCoV in US cattle. In conclusion, we succeeded in the isolation and the comprehensive molecular characterization of a new isolate of BCoV from US cattle. Continuous monitoring of the BCoV genome sequencing is highly recommended for a better understanding of any shift in the viral virulence molecular pathogenesis and to help prepare homologous vaccines against the current circulating strain of the virus in the US cattle population.

## Acknowledgment

We thank Elisa Ramos for her technical assistance with some laboratory activities.

## References

1. Mebus, C.A., et al., Pathology of neonatal calf diarrhea induced by a coronavirus-like agent. Vet Pathol, 1973. 10(1): p. 45–64.

2. McNulty, M.S., et al., Coronavirus infection of the bovine respiratory tract. Vet Microbiol, 1984. 9(5): p. 425–34.

3. Saif, L.J., Bovine respiratory coronavirus. Vet Clin North Am Food Anim Pract, 2010. 26(2): p. 349–64.

4. Rahe, M.C., et al., Bovine coronavirus in the lower respiratory tract of cattle with respiratory disease. J Vet Diagn Invest, 2022. 34(3): p. 482–488.

5. Workman, A.M., et al., Recent Emergence of Bovine Coronavirus Variants with Mutations in the Hemagglutinin-Esterase Receptor Binding Domain in U.S. Cattle. Viruses, 2022. 14(10).

6. Vlasova, A.N. and L.J. Saif, Bovine Coronavirus and the Associated Diseases. Front Vet Sci, 2021. 8: p. 643220.

7. Zhou, Z., Y. Qiu, and X. Ge, The taxonomy, host range and pathogenicity of coronaviruses and other viruses in the Nidovirales order. Anim Dis, 2021. 1(1): p. 5.

8. Workman, A.M., et al., Longitudinal study of humoral immunity to bovine coronavirus, virus shedding, and treatment for bovine respiratory disease in pre-weaned beef calves. BMC Vet Res, 2019. 15(1): p. 161.

9. Zhang, X., et al., Quasispecies of bovine enteric and respiratory coronaviruses based on complete genome sequences and genetic changes after tissue culture adaptation. Virology, 2007. 363(1): p. 1–10.

10. Kiser, J.N. and H.L. Neibergs, Identifying Loci Associated With Bovine Corona Virus Infection and Bovine Respiratory Disease in Dairy and Feedlot Cattle. Front Vet Sci, 2021. 8: p. 679074.

11. Angen, O., et al., Respiratory disease in calves: microbiological investigations on trans-tracheally aspirated bronchoalveolar fluid and acute phase protein response. Vet Microbiol, 2009. 137(1-2): p. 165–71.

12. Hick, P.M., et al., Coronavirus infection in intensively managed cattle with respiratory disease. Aust Vet J, 2012. 90(10): p. 381–6.

13. O’Neill, R., et al., Patterns of detection of respiratory viruses in nasal swabs from calves in Ireland: a retrospective study. Vet Rec, 2014. 175(14): p. 351.

14. Gagea, M.I., et al., Diseases and pathogens associated with mortality in Ontario beef feedlots. J Vet Diagn Invest, 2006. 18(1): p. 18–28.

15. Martin, S.W., et al., The association of titers to bovine coronavirus with treatment for bovine respiratory disease and weight gain in feedlot calves. Can J Vet Res, 1998. 62(4): p. 257–61.

16. Snowder, G.D., et al., Bovine respiratory disease in feedlot cattle: environmental, genetic, and economic factors. J Anim Sci, 2006. 84(8): p. 1999–2008.

17. Jactel B., E.J., Viso M. and Valiergue H., 1990, An Epidemiological Study Of Winter Dysentery In Fifteen Herds In France. Veterinary Research Communications, 1990. 14(5): p. 367-379.

18. Takahashi F., A.H.a.I.Y., Bovine Epizootic Diarrhea Resembling Winter Dysentery Caused by Bovine Coronavirus. Japan Agricultural Research Quarterly, 1983. 17(3): p. 185–190.

19. Takiuchi E., F.B.A., Alfieri A.F., Filippsen P. and Alfieri A.A, An Outbreak of Winter Dysentery Caused by Bovine Coronavirus in a High-Production Dairy Cattle Herd from a Tropical Country. Brazilian Archives of Biology and Technology, 2009. 52: p. 57–61.

20. Toftaker I., H.I., Nødtvedt A., Østerås O., and Stokstad M, A cohort study of the effect of winter dysentery on herd-level milk production. Journal of Dairy Science, 2017. 100: p. 6483–6493.

21. Cho, K.O., et al., Cross-protection studies between respiratory and calf diarrhea and winter dysentery coronavirus strains in calves and RT-PCR and nested PCR for their detection. Arch Virol, 2001. 146(12): p. 2401–19.

22. Gomez, D.E., et al., Detection of Bovine Coronavirus in Healthy and Diarrheic Dairy Calves. J Vet Intern Med, 2017. 31(6): p. 1884–1891.

23. Zhang, M., et al., Respiratory viruses identified in western Canadian beef cattle by metagenomic sequencing and their association with bovine respiratory disease. Transbound Emerg Dis, 2019. 66(3): p. 1379–1386.

24. Health, M.A. Bovine Coronavirus (BCoV) is Highly Prevalent on European Dairy Farms. 2022.

25. Suzuki, T., et al., Genomic Characterization and Phylogenetic Classification of Bovine Coronaviruses Through Whole Genome Sequence Analysis. Viruses, 2020. 12(2).

26. Kim, Y., et al., Trypsin enhances SARS-CoV-2 infection by facilitating viral entry. Arch Virol, 2022. 167(2): p. 441-458.

27. Shin, J., et al., Isolation and Genetic Characterization of a Bovine Coronavirus KBR-1 Strain from Calf Feces in South Korea. Viruses, 2022. 14(11).

28. Reed, L.J. and H. Muench, A SIMPLE METHOD OF ESTIMATING FIFTY PER CENT ENDPOINTS12. American Journal of Epidemiology, 1938. 27(3): p. 493–497.

29. Schneider, C.A., W.S. Rasband, and K.W. Eliceiri, NIH Image to ImageJ: 25 years of image analysis. Nat Methods, 2012. 9(7): p. 671–5.

30. Decaro, N., et al., Detection of bovine coronavirus using a TaqMan-based real-time RT-PCR assay. J Virol Methods, 2008. 151(2): p. 167–171.

31. Livak, K.J. and T.D. Schmittgen, Analysis of relative gene expression data using real-time quantitative PCR and the 2(-Delta Delta C(T)) Method. Methods, 2001. 25(4): p. 402–8.

32. Tamura, K., G. Stecher, and S. Kumar, MEGA11: Molecular Evolutionary Genetics Analysis Version 11. Mol Biol Evol, 2021. 38(7): p. 3022–3027.

33. Pfanzagl, J., Studies in the history of probability and statistics XLIV. A forerunner of the t-distribution. Biometrika, 1996. 83(4): p. 891–898.

34. Mebus, C.A., Infectious enteric viruses of neonatal animals. Am J Clin Nutr, 1977. 30(11): p. 1851–6.

35. Mebus, C.A., L.E. Newman, and E.L. Stair, Jr., Scanning electron, light, and immunofluorescent microscopy of intestine of gnotobiotic calf infected with calf diarrheal coronavirus. Am J Vet Res, 1975. 36(12): p. 1719–25.

36. Salem, E., et al., Global Transmission, Spatial Segregation, and Recombination Determine the Long-Term Evolution and Epidemiology of Bovine Coronaviruses. Viruses, 2020. 12(5).

37. Zhu, Q., B. Li, and D. Sun, Advances in Bovine Coronavirus Epidemiology. Viruses, 2022. 14(5).

38. Hemida, M.G., Middle East Respiratory Syndrome Coronavirus and the One Health concept. PeerJ, 2019. 7: p. e7556.

39. Ko, C.K., et al., Molecular characterization of HE, M, and E genes of winter dysentery bovine coronavirus circulated in Korea during 2002-2003. Virus Genes, 2006. 32(2): p. 129–36.

40. Hasoksuz, M., et al., Molecular analysis of the S1 subunit of the spike glycoprotein of respiratory and enteric bovine coronavirus isolates. Virus Res, 2002. 84(1-2): p. 101–9.

41. Vautherot, J.F., J. Laporte, and P. Boireau, Bovine coronavirus spike glycoprotein: localization of an immunodominant region at the amino-terminal end of S2. J Gen Virol, 1992. 73 **( Pt** **12**): p. 3289–94.

42. Kienzle, T.E., et al., Structure and orientation of expressed bovine coronavirus hemagglutinin-esterase protein. J Virol, 1990. 64(4): p. 1834–8.

43. Lang, Y., et al., Coronavirus hemagglutinin-esterase and spike proteins coevolve for functional balance and optimal virion avidity. Proc Natl Acad Sci U S A, 2020. 117(41): p. 25759–25770.

44. Chang, R.Y. and D.A. Brian, *cis Requirement for N-specific protein sequence in bovine coronavirus defective interfering RNA replication*. J Virol, 1996. 70(4): p. 2201–7.

45. Deregt, D., M. Sabara, and L.A. Babiuk, Structural proteins of bovine coronavirus and their intracellular processing. J Gen Virol, 1987. 68 **( Pt** **11**): p. 2863–77.

46. Bahoussi, A.N., et al., Evolutionary adaptation of bovine coronavirus (BCoV): Screening of natural recombinations across the complete genomes. J Basic Microbiol, 2023. 63(5): p. 519–529.

47. Saif, L.J., et al., Cell culture propagation of bovine coronavirus. J Tissue Cult Methods, 1988. 11(3): p. 139–145.

48. Lin, C.M., et al., Attenuation of an original US porcine epidemic diarrhea virus strain PC22A via serial cell culture passage. Vet Microbiol, 2017. 201: p. 62–71.

49. Li, Y., et al., Structural Requirements and Plasticity of Receptor-Binding Domain in Human Coronavirus Spike. Front Mol Biosci, 2022. 9: p. 930931.

50. Zhang, L., et al., SARS-CoV-2 spike-protein D614G mutation increases virion spike density and infectivity. Nat Commun, 2020. 11(1): p. 6013.

51. Zhang, X.M., et al., Biological and genetic characterization of a hemagglutinating coronavirus isolated from a diarrhoeic child. J Med Virol, 1994. 44(2): p. 152–61.

52. Jeong, J.H., et al., Molecular analysis of S gene of spike glycoprotein of winter dysentery bovine coronavirus circulated in Korea during 2002-2003. Virus Res, 2005. 108(1-2): p. 207–12.

53. Alkan, F., et al., The detection and genetic characterization based on the S1 gene region of BCoVs from respiratory and enteric infections in Turkey. Transbound Emerg Dis, 2011. 58(2): p. 179–85.

54. Hasoksuz, M., et al., Detection of respiratory and enteric shedding of bovine coronaviruses in cattle in Northwestern Turkey. Acta Vet Hung, 2005. 53(1): p. 137–46.

55. Sevinc Temizkan, S. and F. Alkan, Bovine coronavirus infections in Turkey: molecular analysis of the full-length spike gene sequences of viruses from digestive and respiratory infections. Arch Virol, 2021. 166(9): p. 2461–2468.

56. Brandao, P.E., et al., Molecular analysis of Brazilian strains of bovine coronavirus (BCoV) reveals a deletion within the hypervariable region of the S1 subunit of the spike glycoprotein also found in human coronavirus OC43. Arch Virol, 2006. 151(9): p. 1735–48.

57. Bai, Z., et al., The SARS-CoV-2 Nucleocapsid Protein and Its Role in Viral Structure, Biological Functions, and a Potential Target for Drug or Vaccine Mitigation. Viruses, 2021. 13(6).

58. Feng, W., et al., Nucleocapsid protein of SARS-CoV-2 is a potential target for developing new generation of vaccine. J Clin Lab Anal, 2022. 36(6): p. e24479.

59. Huang, Y., et al., Molecular characterization of SARS-CoV-2 nucleocapsid protein. Front Cell Infect Microbiol, 2024. 14: p. 1415885.

60. Jack, A., et al., SARS-CoV-2 nucleocapsid protein forms condensates with viral genomic RNA. PLoS Biol, 2021. 19(10): p. e3001425.

61. Royster, A., et al., SARS-CoV-2 Nucleocapsid Protein Is a Potential Therapeutic Target for Anticoronavirus Drug Discovery. Microbiol Spectr, 2023. 11(3): p. e0118623.

62. Wang, S., et al., Targeting liquid-liquid phase separation of SARS-CoV-2 nucleocapsid protein promotes innate antiviral immunity by elevating MAVS activity. Nat Cell Biol, 2021. 23(7): p. 718–732.

63. Wu, W., et al., The SARS-CoV-2 nucleocapsid protein: its role in the viral life cycle, structure and functions, and use as a potential target in the development of vaccines and diagnostics. Virol J, 2023. 20(1): p. 6.

64. Xiangbo, Z., et al., Bovine coronavirus nucleocapsid suppresses IFN-beta production by inhibiting RIG-I-like receptors pathway in host cells. Arch Microbiol, 2022. 204(8): p. 536.

